# MTSplice predicts effects of genetic variants on tissue-specific splicing

**DOI:** 10.1101/2020.06.07.138453

**Authors:** Jun Cheng, Muhammed Hasan Çelik, Anshul Kundaje, Julien Gagneur

## Abstract

Tissue-specific splicing of exons plays an important role in determining tissue identity. However, computational tools predicting tissue-specific effects of variants on splicing are lacking. To address this issue, we developed MTSplice (Multi-tissue Splicing), a neural network which quantitatively predicts effects of human genetic variants on splicing of cassette exons in 56 tissues. MTSplice combines the state-of-the-art predictor MMSplice, which models constitutive regulatory sequences, with a new neural network which models tissue-specific regulatory sequences. MTSplice outperforms MMSplice on predicting effects associated with naturally occurring genetic variants in most tissues of the GTEx dataset. Furthermore, MTSplice predicts that autism-associated *de novo* mutations are enriched for variants affecting splicing specifically in the brain. MTSplice is provided free of use and open source at the model repository Kipoi. We foresee MTSplice to be useful for functional prediction and prioritization of variants associated with tissue-specific disorders.

## Introduction

Splicing is a fundamental biological process in which introns are cut out from precursor RNAs and exons are joined together. Alternative splicing refers to alternative usage of exons. It is estimated that approximately 95% of human multi-exon genes undergo alternative splicing [1]. Exon skipping (of so-called cassette exons) is the most common alternative splicing pattern [2]. Skipping level of an exon is commonly quantified with the percent spliced-in (PSI or Ψ) [3]. Percent spliced-in can be estimated from RNA-Sequencing (RNA-Seq) data as the number of split RNA-Seq reads supporting the inclusion of the exon divided by the total number of split reads supporting the skipping or the inclusion of the exon. Splicing is a complex process which involves regulation by sequence elements in the exons and flanking introns [4]. Moreover, alternative splicing is often tissue-specific [2, 3, 5, 6]. This means that certain splicing isoforms are only present in certain tissues or that the relative abundances of splice isoforms differ across tissues. Alternative splicing plays an important role in tissue development and shaping tissue identity [7, 8]. Analyzing the protein-coding roles of tissue-specific exons revealed their critical role in rewiring protein interaction networks in different tissues [9]. Tissue-specific splicing patterns are associated with short RNA motifs [2, 10–13]. These short RNA motifs encode tissue-specific splicing regulatory elements, typically intronic or exonic binding sites for splicing factors with a tissue-specific activity. Mammalian tissue-specific splicing factors include Nova1, Nova2, PTB/nPTB, RBFOX1 for nervous tissues, and MBNL1 for muscles, among others. For a review, see Chen & Manley [14].

Splicing defects account for an important fraction of the genetic basis of human diseases [15–17]. Some of these splicing defects are specific to disease-relevant tissues. For instance, individuals affected by autism spectrum disorder (ASD) frequently present mis-splicing of brain-specific exons [18–20] as well as an enrichment of *de novo* mutations in brain-specific exons [21]. Hence, computational tools that can predict the tissue-specific effects of genetic variants on splicing would be relevant for understanding the genetic basis of tissue-specific diseases such as ASD.

Many computational tools have been developed to predict splice sites or splicing strength from sequence [22–32]. However, tools are lacking for predicting tissue-specific effects of human genetic variants on splicing. Barash et al. developed the first sequence-based model predicting tissue-specific splicing in mouse cells [33]. The model integrates regulatory sequence elements to qualitatively predict whether the inclusion of a cassette exon increases, decreases, or remains at a similar level from one tissue to another tissue. This model was further improved to predict directional changes between tissues along with discretized Ψ categories (Low, -Medium, and -High) within a tissue by using a Bayesian neural network with hidden variables [34]. In a subsequent study, a similar Bayesian neural network (SPANR) was trained on human data [28]. However, SPANR was evaluated only for predicting the largest effect across all investigated tissues. Hence, the performance of SPANR on any given tissue is unclear. Moreover, the publicly available SPANR does not allow performing tissue-specific predictions.

We previously developed MMSplice, a neural network with a modular design that predicts the effect of variants on splicing [29, 30]. Unlike SPANR, which has been trained on natural endogenous genomic sequence, MMSplice leverages perturbation data from a recently published massively parallel reporter assay [27]. MMSplice outperformed SPANR and many other splicing predictors in predicting Ψ variations associated with naturally occurring genetic variants as well as effects of variants on percent spliced-in measured on reporter assays [29, 35]. MMSplice models the odds ratio of a cassette exon to be spliced-in when comparing an alternative sequence to a reference sequence.The predicted odds ratios are the same for all tissues because MMSplice has been trained in a tissue-agnostic fashion and therefore does not capture effects of variants affecting tissue-specific regulatory elements.

Deep learning models of tissue-specific regulatory elements have been developed for other biological processes. These models include DeepSEA for chromatin-profiles [36], Basset for DNase I hypersensitivity [37], ExPecto for tissue-specific gene expression [38], FactorNet for transcription factor binding [39], and ChromDragoNN for chromatin accessibility [40]. A common denominator of these models is that they are trained by multi-task learning, i.e. the models make joint predictions for all tissues or cell types using a common set of underlying predictive features. This strategy allows models to efficiently pool information about regulatory elements that are shared across cell types or tissues.

Here, we developed MTSplice (Multi-tissue MMSplice), a model that predicts tissue-specific splicing effects of human genetic variants. MTSplice adjusts the MMSplice predictions with the predictions of TSplice (Tissue-specific Splicing), a novel deep neural network predicting tissue-specific variations of Ψ from sequence which we trained on 56 human tissues using multi-task learning. Performance of MTSplice is demonstrated by predicting tissue-specific variations of Ψ associated with naturally occurring genetic variants of the GTEx dataset as well as investigating brain-specific splicing effect predictions for autism-associated variants. MTSplice is open-source and freely available at the model repository Kipoi [41].

## Results

### Tissue-specific alternatively spliced exons

To train a tissue-specific model of splicing, we considered the alternative splicing catalog of the transcriptome ASCOT [42]. Because the ASCOT annotation and quantification pipeline is annotation-free, it also covers non-annotated exons. Altogether, ASCOT provides Ψ values for 61,823 cassette exons across 56 tissues including 53 tissues from the GTEx dataset [43] and additional RNA-Seq data from peripheral retina. Of note, these tissue-specific values are flagged as missing when the corresponding gene is not expressed [42].

Overall, Ψ of 17,991 exons (29%) of the ASCOT dataset deviate by at least 10% in at least one tissue from its exon-specific average across tissues. These deviations from the exon-specific average Ψ by 10% often occurred in a single tissue (5,658 exons, 31%) and in at least 10 tissues for 4,398 exons (25%, Figure 1A). We investigated co-variations between tissues using these 4,398 exons (Figure 1B). This revealed that samples from the central nervous system (brain, spinal cord, and retina) have very distinct splicing patterns compared to other tissues, in agreement with previous reports [23]. Moreover, skeletal muscle and the two heart tissues (left ventrial and artial appendage) also clustered together with shared splicing patterns. Altogether, this analysis indicates that the ASCOT dataset provides thousands of tissue-specific splicing events that could be used to train a sequence-based predictive model. Also, the ASCOT dataset provides the possibility for a multi-task model to exploit shared splicing regulation of tissues of the central nervous system and, to a lower extent, between skeletal muscle and cardiac tissues.

**Figure 1.**
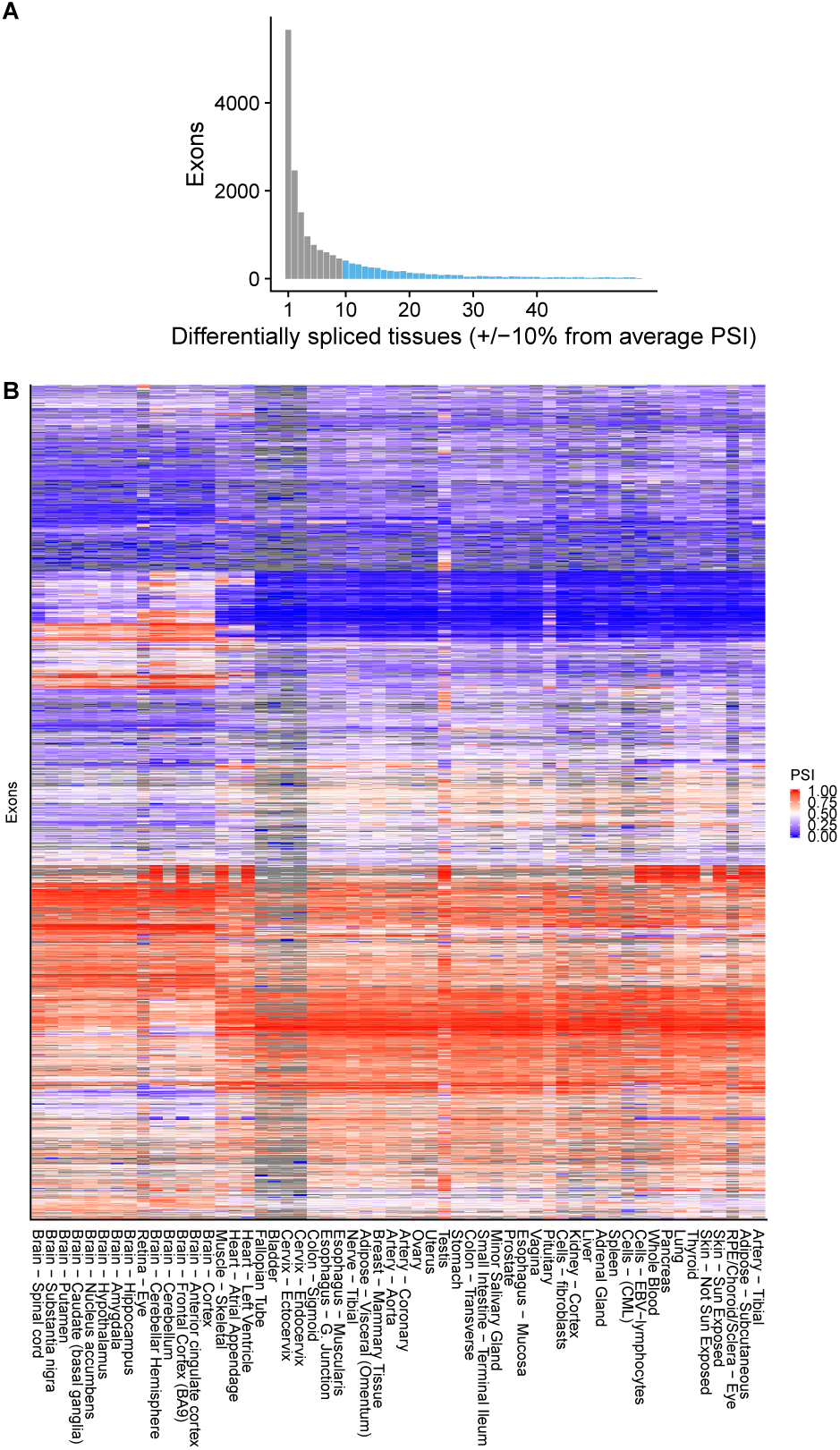
Differential splicing of alternatively spliced exons across tissues. **(A)** Histogram of the number of tissues with differential splicing (Ψ deviating by at least 10% from the exon-average Ψ). Overall, 4,398 exons (light blue) are differentially spliced in at least 10 tissues. **(B)**. Heatmap of Ψ for the 4,398 exons that are differentially spliced in at least 10 tissues with exons (rows) and tissues (columns) sorted by hierarchical clustering. Ψ is color-coded by a gradient from blue (0) to red (1) via white (0.5). Gray entries are missing values and occur in tissues for which the corresponding gene is not expressed. Hierarchical clustering was applied after imputing missing values with row means.

### Differential splicing associated with genetic variants show little tissue-specific variations

The ASCOT dataset consists of data aggregated per tissue. In principle, the genetic variations between donors of the original GTEx dataset provides further information that a sequence-based model could exploit. We therefore next asked how much genetic variation among individuals in GTEx associated with tissue-specific splicing variations. To this end, we computed ΔΨ, the difference between Ψ averaged across individuals homozygous for the alternative allele and Ψ averaged across individuals homozygous for the reference allele for exons with a single variant within the exon body and 300 nucleotides flanking the exon either side (Methods). We estimated Ψ using the software for estimating splice isoform abundances MISO [44], which only takes annotated and alternatively spliced exons into account. Over all these 1,767 single-nucleotide variants, little tissue-specific deviation of ΔΨ compared to its average across tissues was observed (Figure 2A). Specifically, less than 1,476 instances (3.4% of exon-variant-tissue pairs) of tissue-specific ΔΨ deviated by 20% from the tissue-averaged ΔΨ (Figure 2B). This observation is consistent with the fact that only a limited fraction (between 7% and 21%) of splicing QTLs are tissue-specific [45]. Since GTEx samples are derived from healthy donors, this observation, however, does not rule out the possibility that some disease-causing variants do alter splicing in a tissue-specific way. Due to the small amount of tissue-specific splicing variation associated with genetic variants in GTEx, we decided to train a sequence-based model solely based the variations between exons using the ASCOT aggregated data and to keep the genetic variations between donors of the GTEx dataset to independently assess the model afterward.

**Figure 2.**
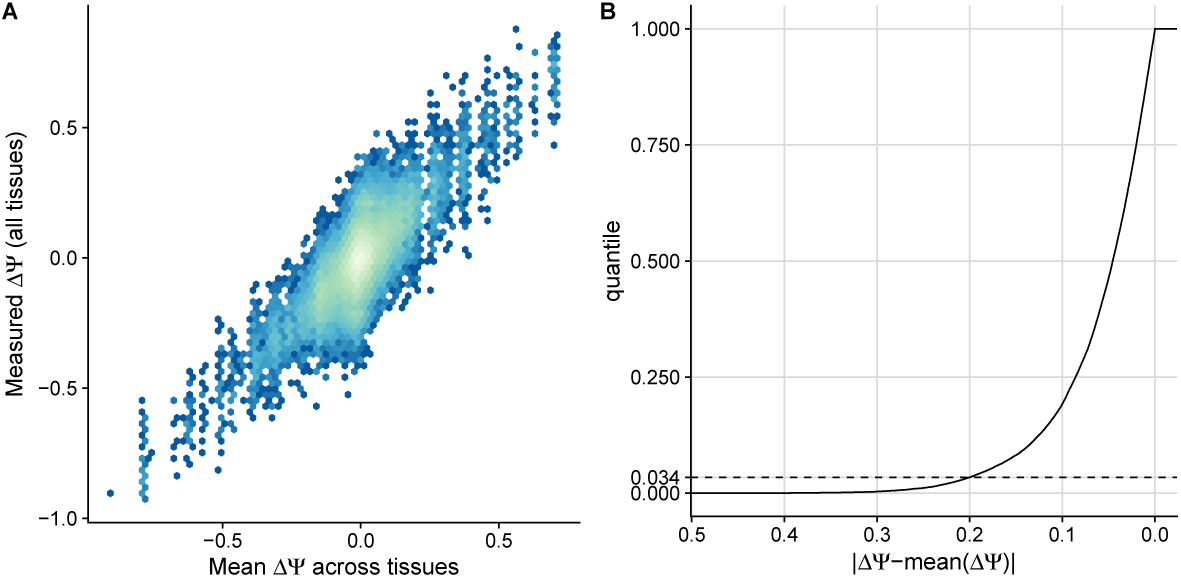
Tissue-specific variations of differential splicing associated with genetic variants in the GTEx dataset. **(A)**. Tissue-specific differential Ψ associated with a genetic variant (y-axis) against differential Ψ associated with a genetic variant averaged across tissues (x-axis). Effects were estimated using homozygous donors **(B)**. Proportion of data points shown in A (y-axis) for every cutoff on the deviation from the averaged Ψ across tissues (x-axis, decreasing).

### TSplice predicts tissue-specific Ψ

We next developed a neural network, TSplice, to predict tissue-specific Ψ values from sequence and tissue-averaged Ψ (Methods). TSplice considers the 300 nt flanking either side of the exon and the first and last 100 nt of the exon body. TSplice is a convolutional neural network (Figure 3) in which positional effects of sequence elements relative to splice sites are modeled using spline transformations [46]. TSplice was trained on the ASCOT dataset using all chromosomes except for chromosome 2, 3 and 5. We report our model prediction performances on these held-out chromosomes.

**Figure 3.**
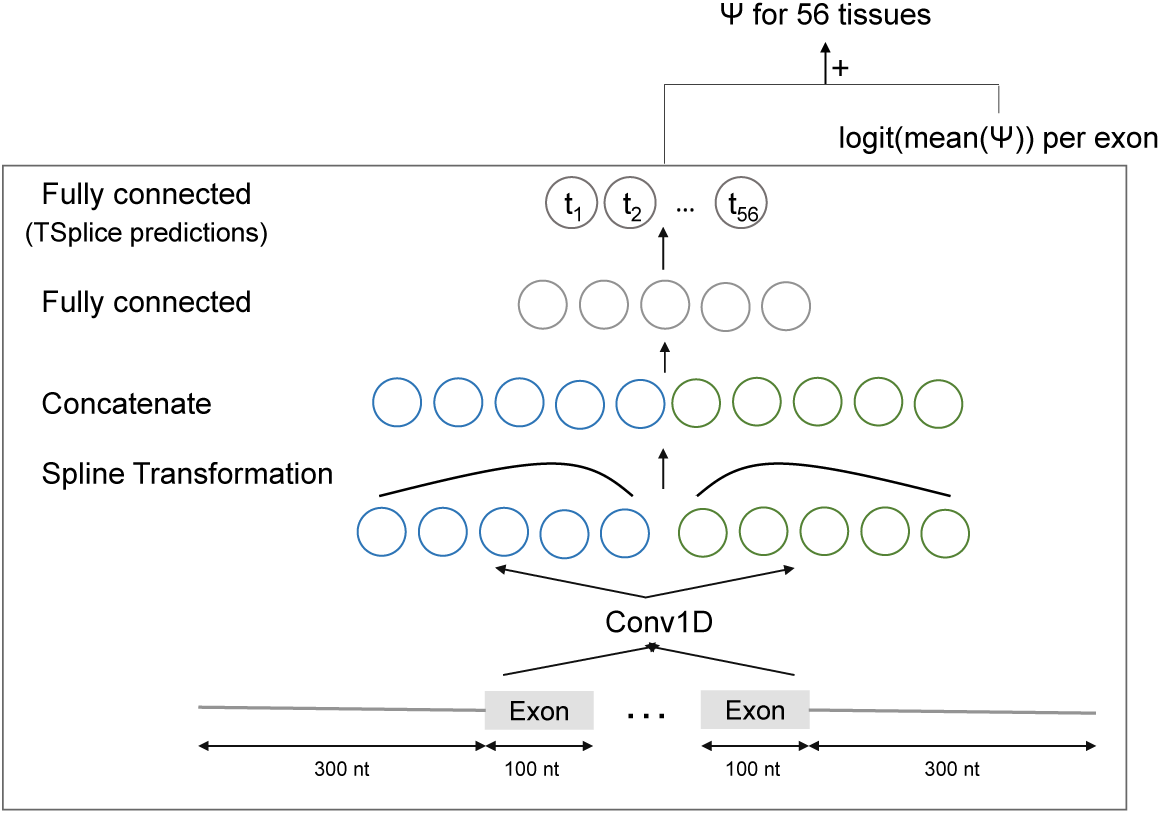
Model architecture to predict tissue-specific percent spliced-in. The model TSplice consists of one convolution layer with 64 length-9 filters capturing sequence elements from one-hot encoded input sequences. This is followed by two spline transformation layers modulating the effect of sequence elements depending on their position relative to the acceptor splice sites (leftmost layer) and the donor (rightmost layer). The outputs of the two spline transformation layers are concatenated and global average pooling is applied along the sequence dimension. This is then followed by feeding two consecutive fully-connected layers. The last fully-connected layer outputs a 56 dimension vector which are the predicted log odds ratios of tissue-specific Ψ versus tissue-averaged Ψ for the 56 tissues of the ASCOT dataset. Natural scale tissue-specific Ψ are obtained by adding predicted odds ratios with measured tissue-averaged Ψ. Batch normalization was used after all layers with trainable parameters except the last fully connected layer. In total, the model has 8,024 trainable parameters.

The performance of TSplice was first assessed on test data by comparing the observed against the predicted log odds ratios of tissue-specific Ψ for 1,621 exons (“variable exons”) with Ψ deviating from the tissue-averaged Ψ by at least 0.2 in at least one tissue and for which the gene is expressed in at least 10 tissues (Figure 4A for Retina eye as an example, Spearman *ρ* =0.27). The predictions positively correlated with the measurements in all tissues and showed a median Spearman correlation of 0.22 (Figure 4B, Supplementary Figure 1). The performance was higher for tissues of the central nervous system (Figure 4C), possibly because central nervous system tissues harbor similar splicing patterns and because they are well represented in the ASCOT dataset.

**Figure 4.**
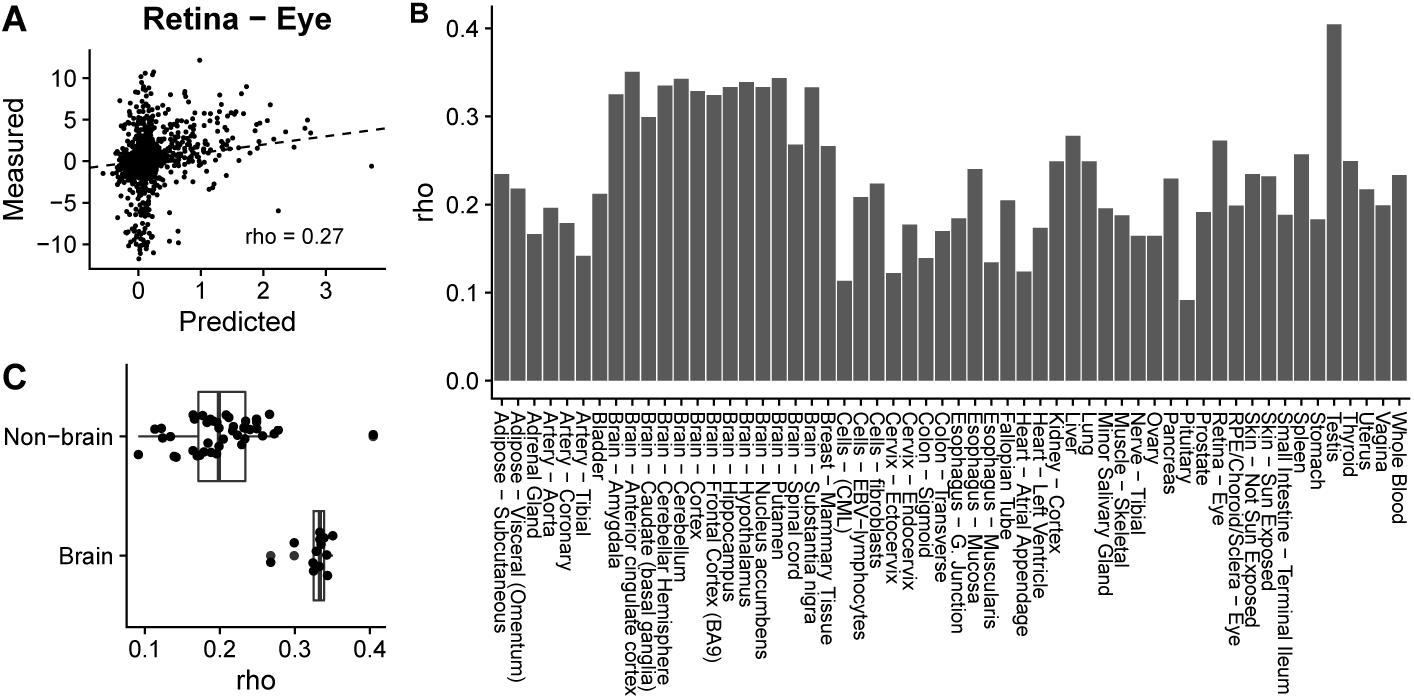
Evaluating TSplice on predicting tissue-associated differential splicing. **(A)** Predicted versus measured tissue-associated differential splicing for the Retina-Eye tissue, representative of the typical performance of our model. **(B)** Spearman correlation between predicted and measured tissue-associated differential splicing for all tissues. **(C)** Distribution of Spearman correlations between predicted and measured tissue-associated differential splicing for brain tissues and non-brain tissues.

We had first assessed log odds ratio predictions because these are the actual quantities the model was trained for. However, percent spliced-ins on the natural scale often matter more for biological and medical applications. We hence next evaluated how well TSplice performs on predicting tissue-specific Ψ on test exons. A successful example is the 9^th^ exon of the gene ABI2, which is included in brain, heart, muscle, and retina tissues and for which TSplice predicts well the order of the tissues (Figure 5A, Spearman *ρ* =0.8) and the absolute values of tissue-specific Ψ per-tissue (root-mean-square error, short RMSE, 0.11). For the majority of the variable exons (73.9%, 1,198 out of 1,621) TSplice ranked tissue-specific Ψ in the right direction (median *ρ* =0.25, Figure 5B). We benchmarked TSplice against a sequence-independent baseline model that predicts tissue-and exon-specific Ψ by adding tissue-average Ψ and exon-average Ψ in the logit scale (Methods). TSplice showed higher Spearman correlation than the baseline model for 65.9% of the exons (*P*< 2.2×10^−16^, paired Wilcoxon test, Supplement Figure S2). Visualization of the positional weights learned by the splines showed that some filters were important for the 5′ half of the model, others for the 3′ half, while about a third of them were important for both halves. Moreover, positional effects were particularly marked near the splice sites (Supplementary Figure S3). To further investigate the motif features the model has learned, we visualized the model gradient on the input sequence. We found the model activates in the presence of known splicing motifs PTBP1, Nova1 and MBNL1 (Supplementary Figure S4). Altogether, these results show that TSplice captured sequence features predictive of Ψ changes across tissues.

**Figure 5.**
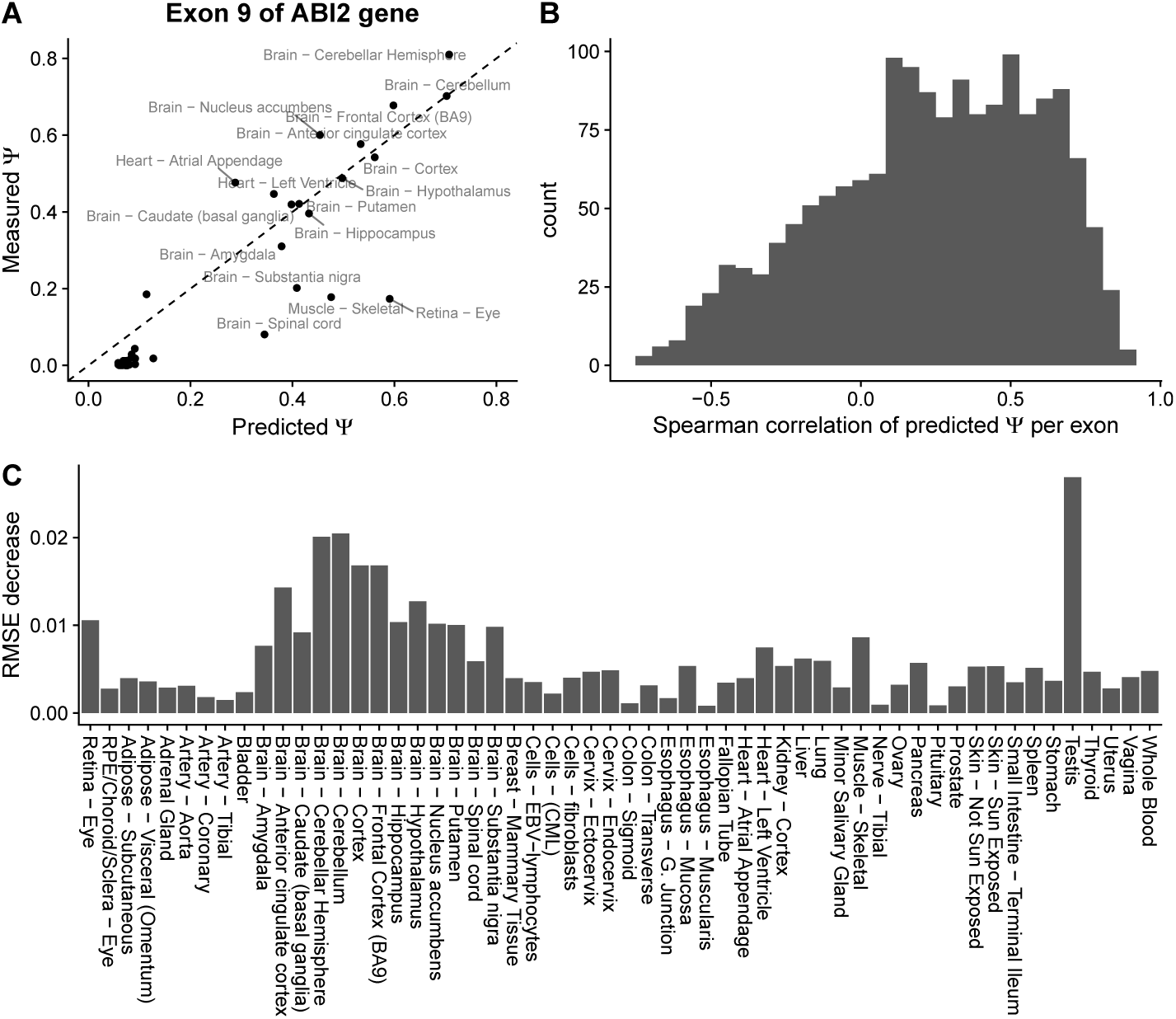
Evaluating TSplice on predicting tissue-specific Ψ. **(A)**. Predicted (x-axis) versus measured (y-axis) Ψ for the 9-th exon of gene ABI2 across 56 tissues. **(B)**. Histogram of the Spearman correlation of the predicted versus measured Ψ for 1,621 test exons across 56 tissues. **(C)** Root-mean-square error decreases between the TSplice model and the baseline model (predicted with the mean Ψ across tissues).

### Tissue-specific variant effect prediction

We next considered combining MMSplice, which models tissue-independent effects together with TSplice, which models differential effects between tissues, to predict the effects associated with genetic variants for any GTEx tissue (Methods). We name this combined model MTSplice. For amygdala, taken as a representative tissue, the MMSplice predictions correlate well (*ρ* =0.42, RMSE=0.188, Figure 6A) with differences of Ψ observed between homozygous donors (Methods). This is consistent with the observation that most variants have similar effects across tissues. Nevertheless, MTSplice further improved the prediction accuracy when evaluated on 1,030 variants with ΔΨ varying by at least 0.2 in at least one tissue (RMSE=0.140 for MMSplice alone, RMSE=0.138 for MTSplice, RMSE=0.141 versus 0.139 when evaluated on all variant). When evaluated on the 51 tissues with at least 10 measured variant effects, MTSplice outperformed MMSplice for 39 out of 51 tissues in terms of root-mean-square error (*P* =1.76 × 10^−5^, paired Wilcoxon test, Figure 6B).

**Figure 6.**
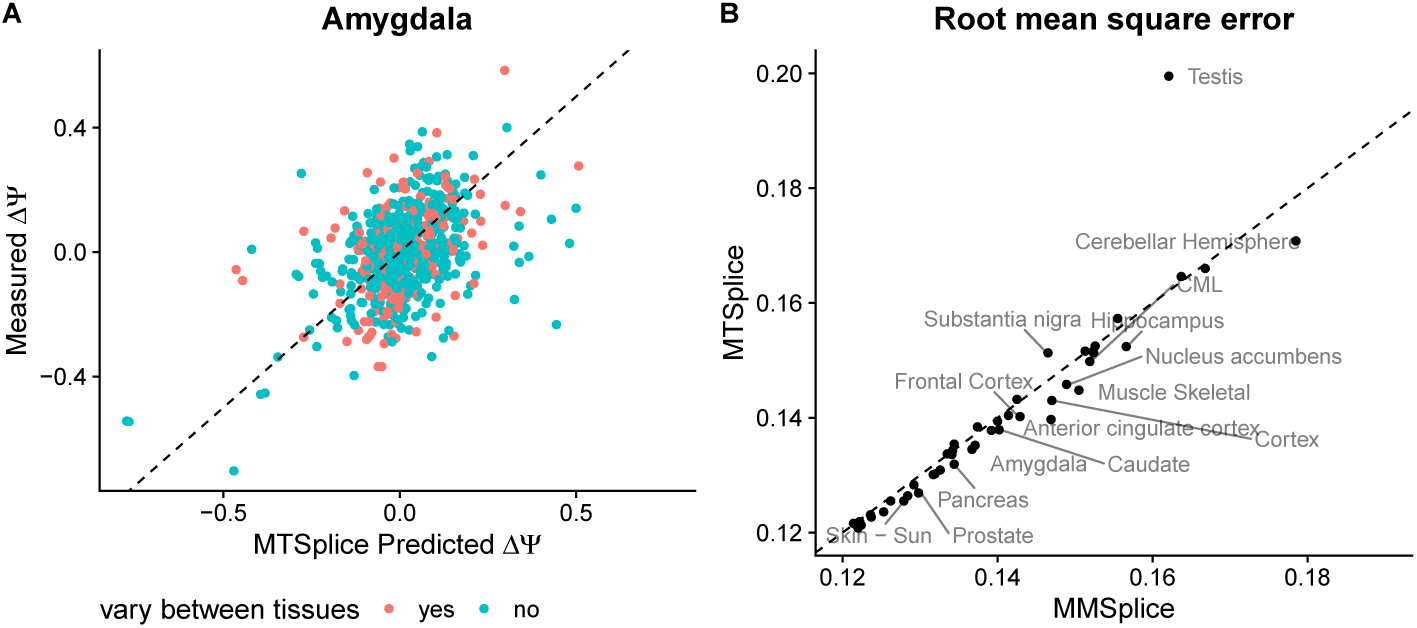
Comparing MMSplice and MTSplice on predicting variant-associated differential splicing. **(A)**. Predicted (x-axis) versus measured (y-axis) ΔΨ in Amygdala between alternative and reference alleles for exons with between-tissue splicing variation (cyan) and other exons (orange). **(B)**. Root-mean-square error of MTSplice predictions (y-axis) against MMSplice predictions (x-asis) for exons with between-tissue splicing variations (the cyan dots in A). Each dot represents one of the 51 GTEx tissues with at least 10 measured variant effects. MTSplice improves for 39 tissues, yet mildly, over MMSplice. Tissues for which the RMSE differences larger than 0.002 are labeled with text.

### MTSplice predicts brain-specific signals for autism patients

To assess the potential of MTSplice on scoring tissue-specific disease variants, we considered *de novo* mutations that were reported for 1,790 autism spectrum disorder (ASD) simplex families from the Simons Simplex Collection [47–51] and as provided by Zhou et al [52]. The data consists of 127,140 *de novo* mutations, with 65,147 from the proband group and 61,993 from the unaffected siblings. Of those, we further considered the 3,884 mutations lying in exons or in their 300 nt flanking intronic regions and predicted with MMSplice with a Δlogit(Ψ) magnitude greater than 0.05. Overall, MMSplice predicted that variants of the proband group would disrupt splicing more strongly than variants of the control siblings (negative MMSplice scores, Figure 7A, *P* =0.042, Wilcoxon rank-sum test). The effect was even stronger for the 1,081 loss-of-function (LoF) intolerant genes (Figure 7A, *P* =0.0035, Wilcoxon rank-sum test, Methods). This result is consistent with the report that LoF-intolerant genes are vulnerable to noncoding disruptive mutations in ASD [52] and points to an important contribution of splicing.

**Figure 7.**
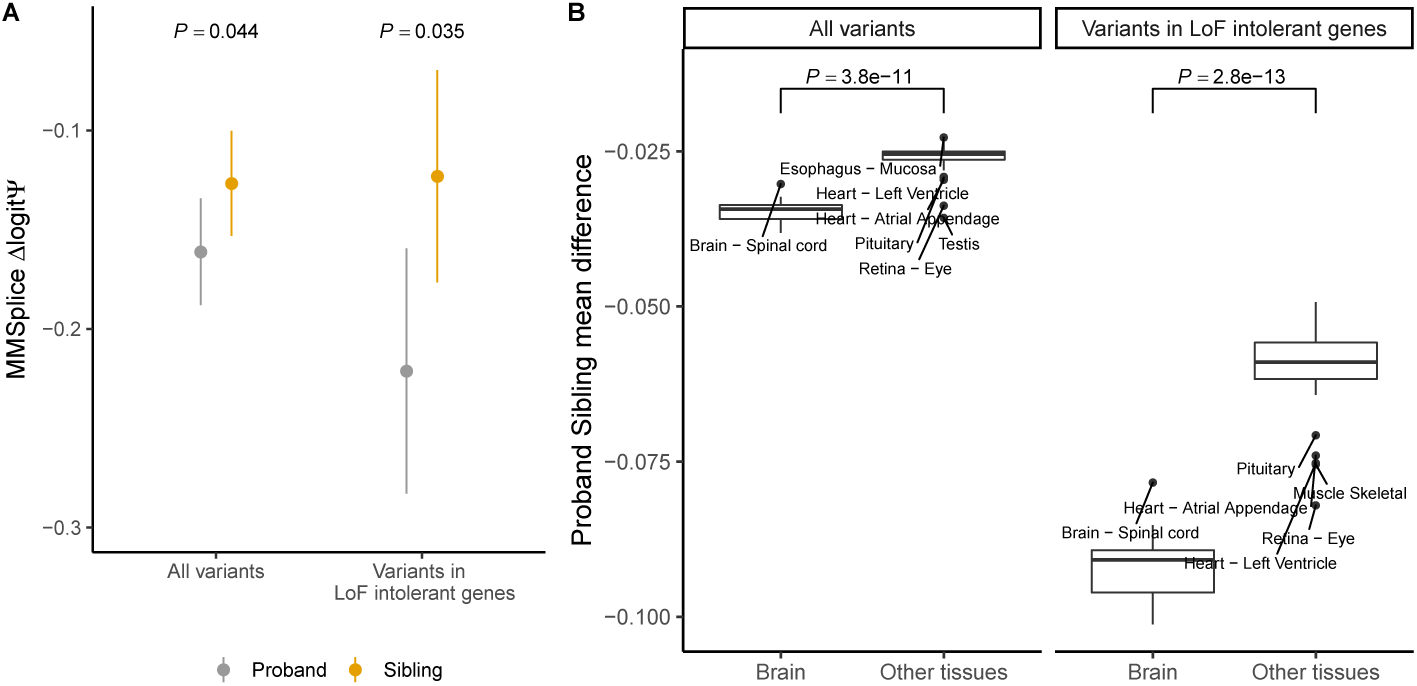
Brain-specific mutational burden on splicing in ASD. **(A)**. Tissue-agnostic variant effect prediction with MMSplice. Splice-region *de novo* mutations (n=3,384, Methods) of the proband group (gray) have significantly lower predicted Δ logit Ψ according to MMSplice compared to those of the unaffected sibling group (orange). The effect size is larger for variants in LoF-intolerant genes (n=1,081). Shown are the means and standard 95% confidence intervals. P-values from one-sided Wilcoxon test. **(B)**. Tissue-specific variant-effect prediction with MTSplice. Distribution of effect size (difference of average Δ logit Ψ for proband versus control siblings *de novo* mutations) for brain tissues (right boxes) and other tissues (left boxes), and for all *de novo* mutations (left panel) or *de novo* mutations in LoF-intolerant genes (right panel) with MTSplice. Individual tissue plots are shown Supplementary Figure S5. The predicted effect sizes are more pronounced for brain tissues.

We then asked whether MTSplice was able to identify tissue-specific effects of ASD-associated *de novo* mutations. Consistent with the MMSplice results, the *de novo* mutations of the proband group were predicted by MTSplice to more severely disrupt splicing than the *de novo* mutations of the control group for all tissues (Figure 7B). The effect size was larger for the brain tissues (Figure 7B). Since autism is a neurological disorder, these results indicate that MTSplice may be used to prioritize variants that could play a tissue-specific pathogenic role. Besides the brain tissues, the tissues with most pronounced differences were retina, which is also part of the central nervous systems and muscle, which has been associated with autism as well [53]. These differences were further amplified when restricting the analysis to the *de novo* mutations in LoF intolerant genes (Figure 7B). Altogether, these analyses demonstrate the value of MTSplice on predicting tissue-specific effects of potentially disease causing mutations.

## Discussion

We introduced the model MTSplice which quantitatively predicts effects of human genetic variants on RNA splicing in 56 tissues. MTSplice has two components. One component, MMSplice, models constitutive splicing regulatory sequences. The other component, TSplice, models tissue-specific splicing regulatory sequences. The combined model MTSplice outperforms MMSplice on predicting tissue-specific variations in percent spliced-in associated with naturally occurring genetic variants in most tissues of the GTEx dataset. Applying MTSplice to *de novo* mutations from autism spectrum disorder simplex families [52], we found a significantly higher burden for the proband group compared to the control siblings, particularly in brain tissues. These results suggest that MTSplice could be applied for scoring variants with a tissue-specific pathogenic role.

The TSplice component was trained from tissue-specific alternative splicing observed in the ASCOT dataset. This approach has two main limitations. First, only less than ten thousand exons show tissue-specific alternative splicing in the ASCOT dataset. This amount of data prohibits training of more complex models. In comparison MMSplice was trained using over 2 million sequences of a massively parallel reporter assay and over half a million naturally occurring splice sites. To overcome this limitation one could leverage complementary data notably tissue-specific expression of splicing-related RNA-binding proteins (RBPs) combined with transcriptome-wide RBP binding profiles [54]. One example of a transfer learning approach in this context is given by Jha et al.[55], who showed the benefits of integrating CLIP-Seq data to predict splicing. The second limitation is that the ASCOT dataset is an observational dataset. Models trained from observational data with genomic sequences may learn sequence features that are correlative but not causal, preventing the models from correctly predicting the effect of genetic variants. This could lead to limited predictive performances of our current model.

One approach to overcome the issue of observational data is to perform massively parallel reporter assays (MPRA) for different cell types. MPRA for human splicing have been performed in HEK293 cells [25, 27, 56–58], K562 cells [59, 60], HepG2 cells [59], and HELA and MCF7 cells [61]. These data provide powerful resources to train complex models on splicing, but tissue and cell-type diversity is still lacking. Tissue-specific MPRA data would also be of prime importance for benchmarking models. Here we had to rely on naturally occurring variants in GTEx for benchmarking. Tissue-specific alteration of splicing can be the outcome of genetic variation affecting either i) constitutive splicing regulatory elements of tissue-specific exons, or ii) tissue-specific splicing regulatory elements. Very few GTEx variants were from the latter class. Hence, the mean square errors differences in GTEx between MTSplice and MMSplice could only be very mild. Previous two-cell-line splicing MPRA experiment did not find tissue-specific variant effects between K562 and HepG2 cells [59], maybe also because the variants tested were selected randomly. A designed MPRA, however, could specifically engineer variations of tissue-specific splicing regulatory elements by using prior knowledge in order to more deeply probe the effect of variants on tissue-specific splicing regulation. The generation of large-scale tissue or cell-type specific perturbation data could therefore be instrumental for probing tissue-specific regulatory elements and could yield more sensitive benchmarks of predictive models.

## Materials and methods

### Dataset

We split the 61,823 cassette exons from ASCOT into a training, a validation, and a test set. The training set consisted of 38,028 exons from chromosome 4, 6, 8, 10-23 and the sex chromosomes. The 11,955 exons from chromosome 1, 7 and 9 were used as the validation set, and the remaining 11,840 exons were used as the test set (chromosomes 2, 3 and 5). Models are evaluated based on their performance on the test set.

### Variant effect estimation

To compute variant effect, we first computed Ψ with MISO for all annotated alternatively spliced exons (MISO annotation v2.0, http://genes.mit.edu/burgelab/miso/annotations/ver2/miso_annotations_hg19_v2.zip) in all GTEx RNA-Seq samples. This led to Ψ estimates for 4,686 samples from 53 tissues. Second, for each exon, we estimated variant effects using only those samples with a single variant within the exon body and 300 nt flanking of the exon. Third, we estimated the effect associated with the variants as the difference between Ψ averaged across samples homozygous for the alternative allele and Ψ averaged across samples homozygous for the reference allele. We required at least 2 samples in each of these two groups. For simplicity, we did not consider heterozygous samples for estimating the effects because Ψ of heterozygous samples is confounded by allele-specific RNA expression. Also, we did not consider indels.

### The TSplice model

We denote Ψ_*e,t*_ the percent spliced-in value of the cassette exon *e* in tissue *t*. The goal of the multi-tissue splicing model is to predict tissue-specific Ψ_*e,t*_ from the nucleotide sequence of the given exon S_*e*_. We train the tissue-specific splicing model with multi-task learning, where each task corresponds to a tissue. The model has two input branches. The first input branch consists of the sequence 300 nt upstream of the acceptor and 100 nt downstream of the acceptor (Figure 3). In a symmetric fashion, the second input branch consists of the sequence from the donor side, with 100 nt upstream of the donor and 300 nt downstream of the donor. All input sequences are one-hot encoded. The input layer is followed by a 1D convolution layer with 64 filters of length 9. Parameters of the convolution layer are shared by the two input branches, based on the assumption that many sequence motifs are presented both upstream and downstream of the exons. To model the positional effects of splicing motifs, spline transformations [46] are fitted for each of the convolution filters to weight the convolution activations based on the relative input position to donor and acceptor sites. The spline transformations are fitted differently for the two input branches to account for potential different positional effects of the upstream and downstream introns. The weighted activations are then concatenated along the sequence dimension. Two fully-connected layers are followed after the concatenated outputs. The last fully-connected layer output number of predictions equals the number of tissues (*T*), corresponding to predictions for each tissue. These are the predictions of the TSplice model mentioned in the manuscript. During training, logit of the mean Ψ per exon 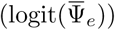 was added to these prediction outputs followed by a sigmoid function. This encourages the model to learn sequence features associated with differential splicing across tissues.

Formally, for each exon, TSplice predicts for each tissue its Ψ_e,t_ deviation from the mean 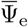 across tissues on logit level. Specifically, we define the tissue-associated differential splicing as Δ_tissue_ logit(Ψ_*e,t*_)

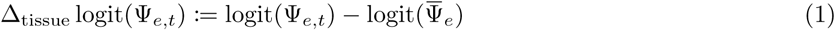

as the logit Ψ deviation for tissue *t* and exon *e* from the logit of 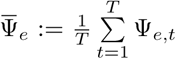 the mean Ψ across tissues.

For exon *e* with input sequence S_*e*_, TSplice predicts the target in ℝ^*T*^ : TSplice(S_*e*_) := (Δ_tissue_ logit(Ψ_*e*,1_), …, Δ_tissue_ logit(Ψ_*e,T*_)) corresponding to *T* tissues.

The tissue-specific Ψ_*e,t*_ can be predicted with TSplice and the given 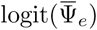 computed from the data as:

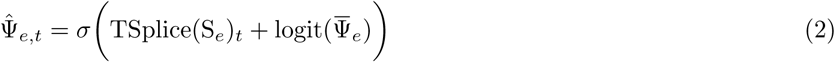

where TSplice(S_*e*_)_*t*_ is the TSplice predicted Δ_tissue_ logit(Ψ_*e,t*_), and *σ* is the sigmoid function: 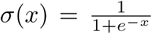. Note that in Equation 1 and elsewhere the average was computed before and not after logit-transformation because it gave more robust results.

### Model training and selection

The model was implemented with keras (version 2.2.4). The Kullback–Leibler (KL) divergence between the predicted and measured Ψ distribution was used as the loss function (Equation 3), by considering the percent spliced-in as the probability of the cassette exon to be included in any given transcript.

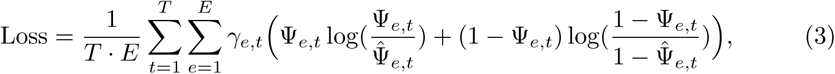

where

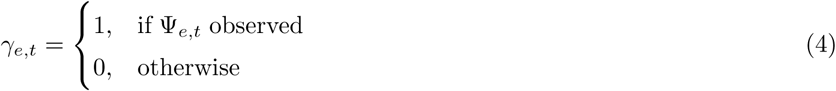

Missing values, which typically correspond to tissues in which the gene is not expressed, were masked out in the loss function. Ψ values were clipped to be between [10^−5^, 1 -10^−5^]. Adam optimizer [62] with default parameters was used to optimize the model. Network weights were initialized with the He Normal initialization [63]. Hyperparameter search was performed with hyperopt [64] with the Tree Parzen Estimators method along with the package kopt (https://github.com/Avsecz/kopt). Hyperparameters were selected based on the loss on the validation set.

After finding the best hyperparameter combination, 20 models were trained with the best hyperparameters but different random initialization. A forward model selection strategy was used to select a set of models whose average predictions gives the smallest loss on the validation set. To this end, models were first sorted based on their performance on the validation set. Next, models were successively added to an ensemble model, defined as the average over the selected models, until the validation set performance no longer improved. This procedure yielded an ensemble model composed of 8 individual models. TSplice predictions are made by this ensemble model.

### Baseline tissue-specific Ψ_*e,t*_ prediction model

The following model was considered as a baseline model to predict tissue-specific 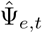:

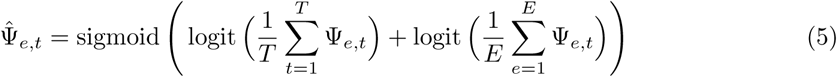

As for the TSplice model, logit-transformation was performed after averaging rather than the other way round because it gave more robust results.

### Tissue-specific variant effect prediction

Tissue-specific variant effect ΔΨ_*e,t*_ is predicted as follows (we considered in this study only homozygous cases as described in the Methods subsection “Variant effect estimation”):

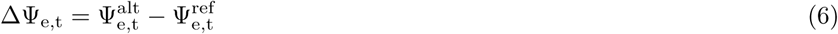

where 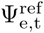 is the measured Ψ for exon e and tissue *t* with the reference sequence, and 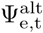 is the tissue-specific Ψ with the alternative sequence. We model the logit level of 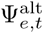 with the following linear model:

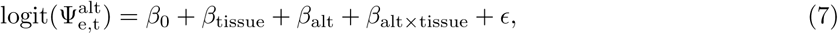

where *β*_0_ is intercept, *β*_tissue_ is the tissue effect, *β*_alt_ is the effect of the variant on an average tissue, *β*_alt×tissue_ is the interaction term which we model the interaction of the variant effect and the given tissue. We model each of the terms as follow:

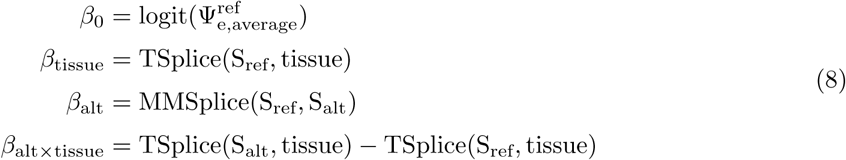

When we plug in Equation 8 into Equation 7, we obtain the MTSplice model which combines MMSplice and TSplice to model tissue-specific variant effect:

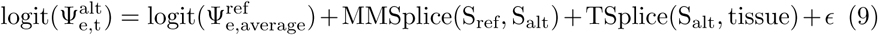

Finally, the tissue specific ΔΨ_e,t_ is predicted as follow:

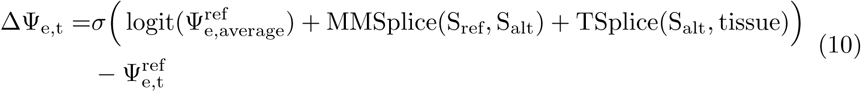

### Benchmark variant effect prediction on GTEx

On the benchmark of tissue-specific variant effect prediction, we further applied four filters. First, we selected variants that have |ΔΨ_*e,t*_| > 0.2 in at least one tissue. Second ΔΨ can be computed for at least 3 tissues. Third, we only considered tissues with more than 10 variants satisfying the above criteria. Altogether, these filters led to 1,030 variant-exon pairs and 51 tissues used for benchmarking tissue-specific variant effect predictions.

### Autism variants

The processed *de novo* mutations were downloaded from the link provided by Zhou et al (Zhou et al. 2019) (https://hb.flatironinstitute.org/asdbrowser/). The original whole genome sequencing data were accessed through the Simons Foundation Autism Research Initiative (SFARI) [47–51]. The data provides 127,140 single nucleotide variants (SNVs) from non-repeat-region. The variants were derived from 7,097 whole genomes from the Simons Simplex Collection (SSC) cohort, which consists of whole-genome sequencing data from 1,790 families (with probands and matched unaffected siblings).

To predict variant effect on splicing, variants were mapped to exons if they are within the annotated (ensembl gene annotation v75) exon body or within 300 nt flanking. If a variant was mapped to multiple exons, the largest effect size was reported as the effect of the variant. A total of 13,415 variants were mapped to known exons and therefore were predicted by our models. Among those variants, 3,884 have predicted |Δlogit(Ψ)| > 0.05. We classified the variants into loss-of-function (LoF) group and loss tolerant group based on the loss-of-function observed/expected (oe) upper bound fraction (LOEUF) scores [65]. We used the suggested cutoff of 0.35 on the upper bound of the oe confidence interval to group the variants.

## Author’s contributions

JC and JG designed the model with the help of AK. JC and MHÇ implemented the software. JC performed the analysis. JG and AK supervised the project. JC and JG wrote the manuscript.

## Acknowledgements

We thank Nils Wagner, Florian Hölzlwimmer and Leonhard Wachutka for their feedback on the manuscript.

## Availability

MTSplice is integrated with MMSplice and is available from https://github.com/gagneurlab/MMSplice. Analysis code is available from https://gitlab.cmm.in.tum.de/gagneurlab/mtsplice

## Additional Files

Additional file 1 — Supplemental Figures

Supplemental Figures S1-S5.

