## Supplement Figures for "MTSplice predicts effects of genetic variants on tissue-specific splicing"

### 1 Supplementary Figures

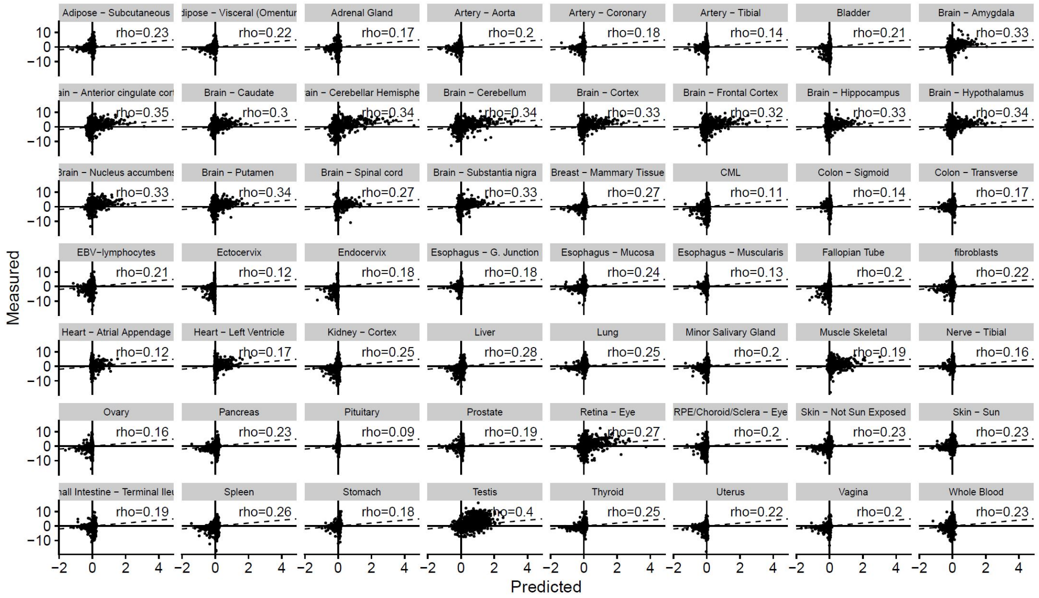


**Supplement Figure S1**: Predicted (x-axis) versus measured (y-axis) tissue-associated differential splicing. The dashed line indicates the x=y line.

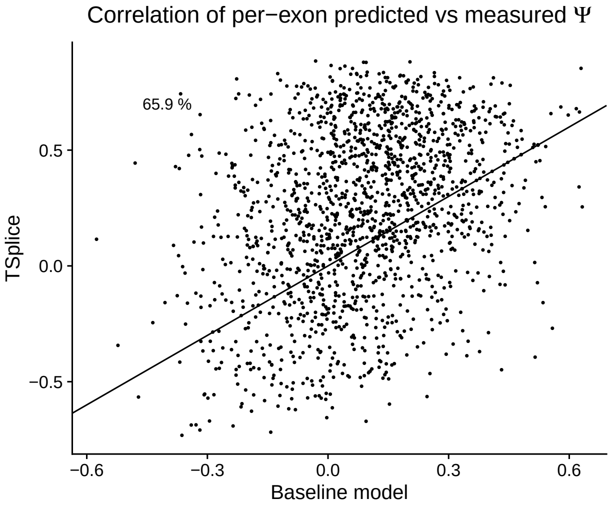


**Supplement Figure S2**: Comparing the spearman correlation between the predicted and the measured tissue-specific $\Psi_{e,t}$ per exon across tissues with a baseline model predicting tissue-specific $\Psi_{e,t}$ by adding the mean $\Delta\Psi$ across exons to the mean $\Psi_e$ across tissues (Methods). TSplice outperforms the baseline model for 64.5 % of the evaluated exons. All exons in the test set with $\Psi$ vary ($|\Psi_{e,t} - \Psi_e| > 0.2$) in at least one tissue and is expressed in at least 10 tissues are considered (1621 exons).

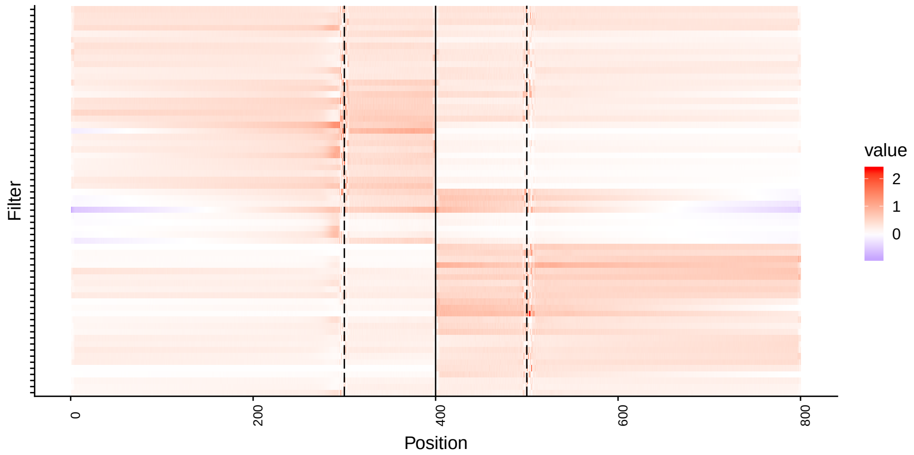


**Supplement Figure S3**: Average activation map of the convolution layer weighted by the spline transformation layer. Dashed lines represent exon-intron boundary.

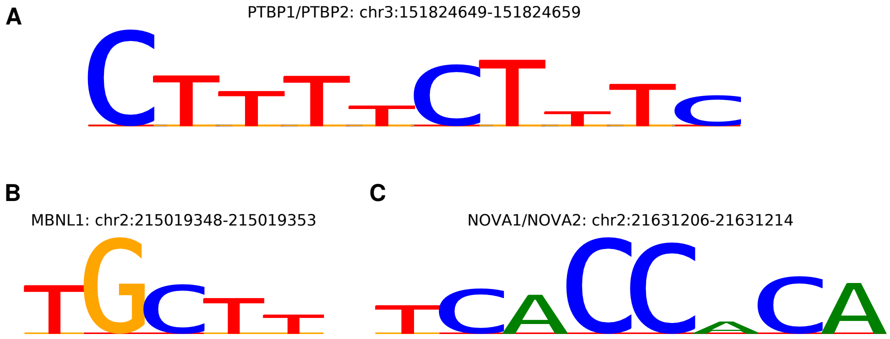


**Supplement Figure S4**: Motif instances detected by TSplice model.

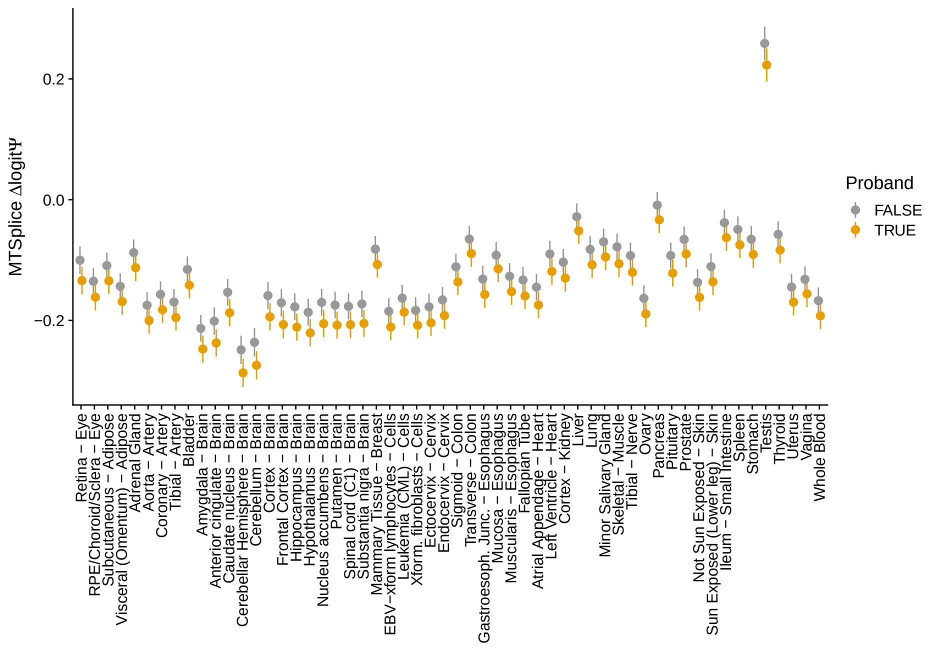


**Supplement Figure S5**: Tissue-specific variant-effect prediction with MT-Splice. MTSplice predicted variant-effect ($\Delta\,\mathrm{logit}\,\Psi$, y-axis) for proband (gray) and the unaffected sibling group (orange) across 56 tissues. The means and standard 95% confidence intervals were shown. P-values from one-sided Wilcoxon test.
